# Conservation and Divergence of YODA MAPKKK Function in Regulation of Grass Epidermal Patterning

**DOI:** 10.1101/287433

**Authors:** Emily Abrash, M Ximena Anleu Gil, Juliana L Matos, Dominique C Bergmann

## Abstract

All multicellular organisms must properly pattern cell types to generate functional tissues and organs. The organized and predictable cell lineages of the *Brachypodium* leaf enabled us to characterize the role of the MAPK kinase kinase gene *BdYODA1* in regulating asymmetric cell divisions. We find that *YODA* genes promote normal stomatal spacing patterns in both *Arabidopsis* and *Brachypodium*, despite species-specific differences in those patterns. Using lineage tracing and cell fate markers, we show that, unexpectedly, patterning defects in *bdyoda1* mutants do not arise from faulty physical asymmetry in cell divisions but rather from improper enforcement of alternative cellular fates after division. These cross-species comparisons allow us to refine our interpretations of MAPK activities during plant asymmetric cell divisions.

**Summary Statement:** Analysis of *Brachypodium* leaf epidermis development reveals that the MAPKKK, *BdYODA1*, regulates asymmetric divisions by enforcing resultant cell fates rather than driving initial physical asymmetries.

## Introduction

The correct establishment of cell types during development is essential for the generation of cellular diversity and patterning of tissues, organs, and organisms. Asymmetric cell division, the process that gives rise to daughter cells with different physical appearance and/or developmental fate, is a crucial mechanism that most eukaryotes employ to generate diverse but organized cell populations. Asymmetric cell divisions are necessary from the very first to the very final events in plant development, and have been studied extensively during embryogenesis, root, shoot, and reproductive development in flowering plants (Van Norman, 2016; Marzec et al., 2015; Abrash and Bergmann, 2009) as well as in basal lineages (Harrison et al., 2009; De Smet and Beeckman, 2011), albeit in a more limited fashion. A number of regulatory players and mechanisms have been identified in specific cellular contexts; however, unifying players and modules used repeatedly among these different asymmetric cell division contexts are only just beginning to come to light (Abrash and Bergmann, 2009; De Smet and Beeckman, 2011).

The stomatal lineages of various flowering plants offer particularly rich and accessible model systems for the study of asymmetric division regulation. Stomata are epidermal valves consisting of paired guard cells flanking a central pore, and they contract and relax to regulate gas exchange between the plant and its environment. In many plant species, asymmetric cell divisions produce one daughter cell that acts as a stomatal precursor and another that differentiates as an epidermal pavement cell (Fryns-Claessens and Van Cotthem, 1973; Dong and Bergmann, 2010; Lau and Bergmann, 2012). In broadleaf dicots like *Arabidopsis*, the asymmetric divisions are self-renewing–that is once a cell undergoes an asymmetric division, its progeny may also continue dividing asymmetrically, in a situation somewhat analogous to the transit amplifying cells in animal lineages (Matos and Bergmann, 2014). In the grasses, by contrast, the stomatal lineage exhibits a less flexible pattern of divisions in which stomatal precursors undergo a single asymmetric division that yields a terminal precursor (the guard mother cell, or GMC) and a larger sister cell fated to become a pavement cell. Also, notably in grasses, the lateral neighbors of the GMC undergo unique additional asymmetric divisions, parallel to the long axis of the leaf, to form the subsidiary cells that act to facilitate stomatal function (Hepworth et al., 2017). In *Brachypodium distachyon*, a forage grass related to wheat, asymmetric cell divisions accompany the creation of all characterized epidermal cell types (Raissig et al., 2016). Here, both stomatal and hair cell lineage development are subject to tight temporal regulation and occur in a developmental progression from the base (younger tissue) to the tip (older tissue) of the leaf blade (Fig. 1A-B). The common deployment of asymmetric divisions in multiple epidermal lineages in grasses requires that stomatal fate be later superimposed in particular cell files to specify the smaller daughter cells as stomatal precursors (Raissig et al., 2016).

**Figure 1.**
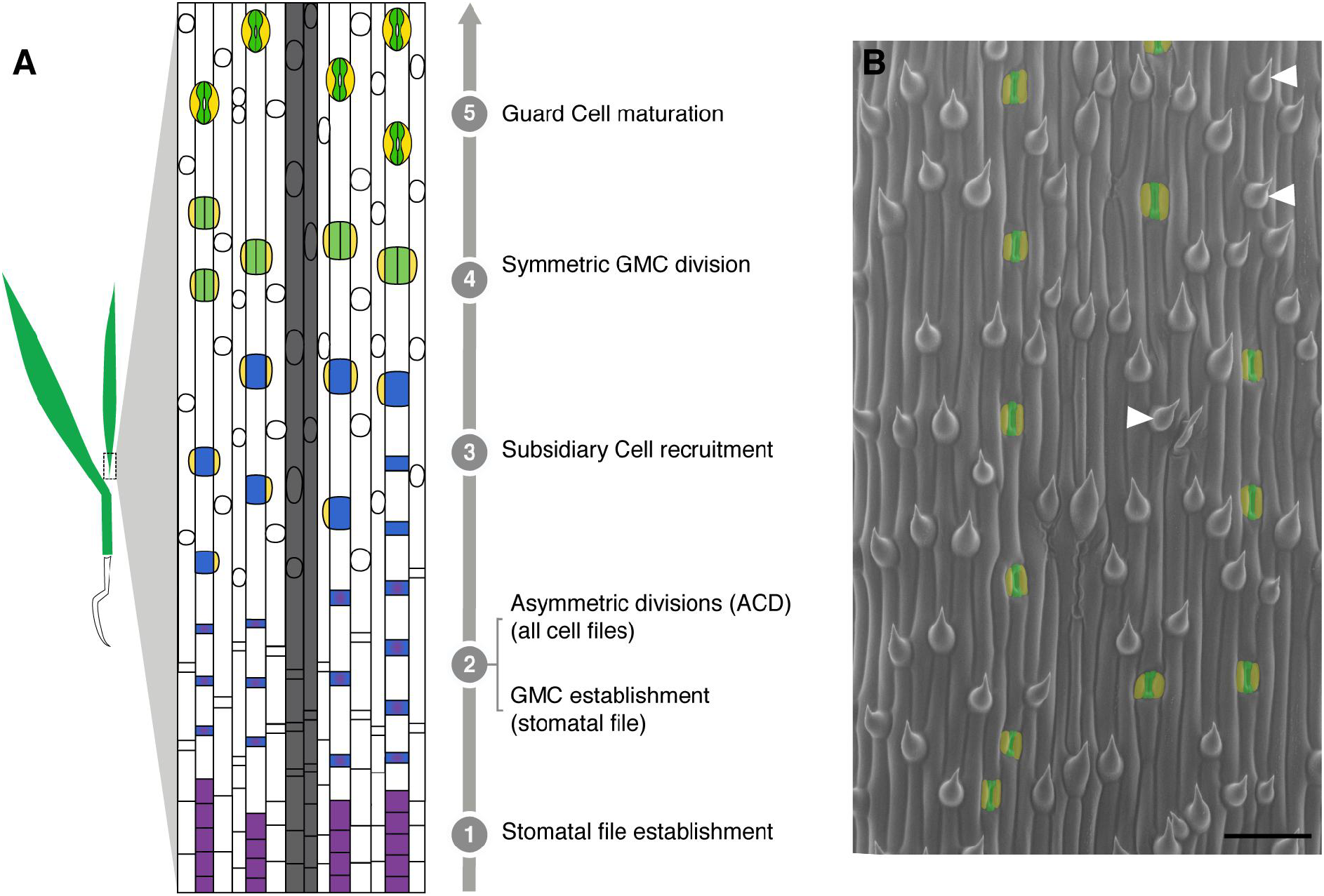
Stomatal development in *Brachypodium* as model to study the progression of asymmetric divisions. **A)** Simplified model of leaf blade epidermal development in *Brachypodium.* Specific cell files at predictable distances from veins (grey files) acquire stomatal lineage fate (stage 1) and undergo stomatal differentiation in a tip-to-base gradient. All cells in the epidermis then divide asymmetrically (ACD). In stomatal files, the smaller daughter cell of each division becomes a Guard Mother Cell (GMC, blue, stage 2). In all other files, these cells develop into hair cells (white circles in non-stomatal files). GMCs then recruit Subsidiary Cells (SC, yellow, stage 3), divide once symmetrically to form two guard cells (green, stage 4), and mature as 4-celled complexes (stage 5). **B)** Scanning electron micrograph of WT (Bd21-3) leaf epidermis. Guard cells and subsidiary cells are false-colored green and yellow, respectively. Co-existence of stomatal and hair fates in a single file is highlighted by white arrowheads. Scale bar = 50 µm.

Asymmetric divisions during development are guided by fate, polarity, and signaling inputs. In *Arabidopsis* stomatal production, bHLH transcription factors, the polarity protein BASL, and a signaling pathway comprising ligands, receptors, and a mitogen-activated protein kinase (MAPK) cascade have been connected to these roles (Lau and Bergmann, 2012; Pillitteri et al., 2016). The MAPK pathway is headed by the MAPK kinase kinase (MAPKKK) YODA (AtYDA) (Bergmann et al., 2004). Genetic and biochemical data indicate that AtYDA responds to positional information provided by peptide ligands of the EPIDERMAL PATTERING FACTOR (EPF) family via transmembrane receptors TOO MANY MOUTHS (TMM) and members of the ERECTA (ER) family (ERf) (Nadeau and Sack, 2002; Kim et al., 2012; Hunt and Gray, 2009; Kondo et al., 2010; Sugano et al., 2010; Hara et al., 2007; Shpak et al., 2005). AtYDA then relays this information through downstream kinases MKK4/5 and MAPK3/6 (Lampard et al., 2009; Wang et al., 2007) to phosphorylate and inhibit the bHLH transcription factor SPEECHLESS (AtSPCH), the primary regulator of entry into the stomatal lineage pathway (Lampard et al., 2008). Loss of *AtYDA* activity results in the production of excess stomata arranged in large clusters, whereas overactivity suppresses development of stomata (Bergmann et al., 2004), phenotypes opposite of those ascribed to loss and gain of *AtSPCH* function (Lampard et al., 2009; Macalister et al., 2007).

In grasses, the stomatal lineage requires homologues of the bHLH transcription factors known from *Arabidopsis*, though sometimes employed in different ways (Raissig et al., 2016; Raissig et al., 2017; Liu et al., 2009). *BdICE1*, *BdSPCH1*, and *BdSPCH2* are needed for stomatal lineage initiation, and the two *SPCH* homologues are expressed in epidermal cells before they undergo asymmetric cell divisions, but their expression becomes restricted to the smaller daughter cells after division (Raissig et al., 2016). EPF overexpression was shown to arrest stomatal production in barley (Hughes et al., 2017), potentially acting upstream of these transcription factors, but whether EPFs enforce stereotyped asymmetric divisions and the “every other cell” epidermal pattern in grasses is yet unknown.

In a screen aimed at identifying mutations that affect stomatal patterning in *Brachypodium*, we identified a mutation in *BdYDA1* that displays clustered stomata similar to loss of *AtYDA* in *Arabidopsis*. The mutation in *BdYDA1* also led to patterning and fate defects in other epidermal cell types and changes in overall plant morphology. These additional phenotypes were interesting in light of previous findings that *AtYDA* is not exclusively a stomatal lineage regulator–it was originally characterized for its role in asymmetric divisions of the zygote (Lukowitz et al., 2004) and was subsequently shown to act in the development of the inflorescence (Meng et al., 2012) and anthers (Hord et al., 2008), in defense (Sopeña-Torres et al., 2018), and in specifying division plane orientation in the root (Smékalová et al., 2014). Based on work in *Arabidopsis*, YDA was surmised to establish physically asymmetric divisions that lead to different cell fates in the daughters. By taking advantage of the highly spatially and temporally organized development of the *Brachypodium* leaf, however, we show that patterning defects do not arise from a fault in the physical asymmetry of cell divisions, but from improper enforcement of alternative cellular fates in these tissues. This comparative work supports a role for YDA as a conserved regulator of asymmetric cell divisions, but expands our understanding of its role within those divisions.

## Results

### Mutations in *BdYDA1* lead to severe disruptions in stomatal pattern

In wild-type *Brachypodium distachyon* Bd21-3 (WT), stomata are separated by at least one intervening non-stomatal cell (Fig. 1A-B). From an EMS-mutagenized population of WT plants (described in Raissig et al., 2016), we identified a line segregating plants that, unlike WT, bore large groups of stomata in contact in a single row (Fig. 2A-B, H). The mutant did not exhibit ectopic stomatal rows, suggesting that the mutation affected processes occurring after specification of stomatal row identity. In addition to the stomatal patterning defects, mutants displayed whole plant growth defects, including compressed internodes and lateral organs, reduced overall stature, dark green coloration, and sterility (Fig. 2E). This combination of phenotypes was similar to that previously described for *Arabidopsis* plants bearing mutations in the MAPKKK encoding gene, *AtYDA* (Bergmann et al., 2004; Lukowitz et al., 2004). *YDA* orthologues can be identified in many plant species; in the grasses, *YDA* has been duplicated (Fig. S1). We sequenced the gene with the higher sequence similarity to *AtYDA* [*BdYDA1* (Bradi5g18180)] and found a missense mutation (G2287A) predicted to confer a charge change (E460>K) in a conserved residue of the kinase domain (Fig. 2I).

**Figure 2.**
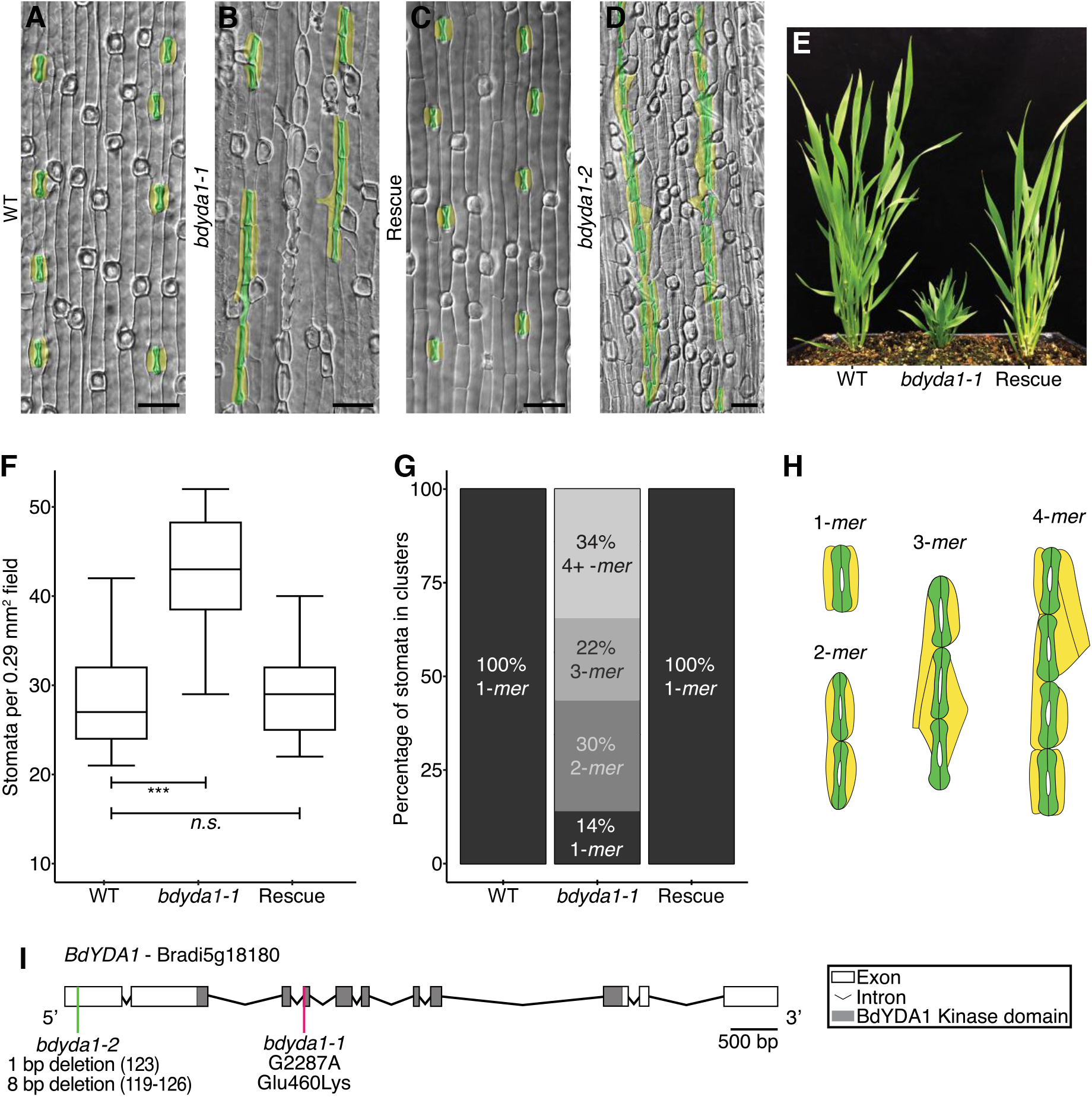
*BdYDA1* is required for proper spacing of stomata. **(A-D)** Differential Interference Contrast (DIC) images of cleared WT (Bd21-3) (A), *bdyda1-1* mutant (B), *bdyda1-1* complemented with *BdYDA1pro:BdYDA1-YFP:Yt* transgene (C), and *bdyda1-2* (D) abaxial leaf epidermis. Guard cells and subsidiary cells are false-colored green and yellow, respectively. Images are of the 6^th^ leaf from base (3^rd^ from main tiller) 27 days post-germination (dpg) with WT and rescued line images being cleared leaves and *bdyda1-1* images from epidermal peels. *bdyda1-2* image shows cleared leaf from T0 regenerant. Scale bars = 40 µm. **(E)** Whole-plant phenotype of *bdyda1-1* mutant (middle), WT (right), and rescue (5 weeks post germination). **(F)** Stomatal density of *bdyda1-1* mutants compared to that of WT and rescued *bdyda1-1* [6^th^ leaf from base (3^rd^ from main tiller) 27 dpg]. n=4 individuals for WT control and n=5 for rescued plants. For each sample, 5 different regions of the leaf were imaged and quantified. n=5 for *bdyda1-1* mutants for which 4 different regions of the leaf were peeled, imaged, and quantified. ***, significant at p<0.001; *n.s*., not significant (based on Kruskal-Wallis test followed by Dunn’s multiple comparison’s test). In boxplot, the black horizontal line indicates the median; hinges (upper and lower edges of the box) represent versions of the upper and lower quartiles; whiskers extend to the largest observation within 1.5 interquartile ranges of the box. **(G)** Stomatal cluster profile as percentage of clustering of quantified stomata in (F) (n=566 stomata for WT controls, n=729 stomata for rescue, and n=835 stomata for *bdyda1-1*). Clusters of 4 or more stomata were grouped in last category “4+ *-mer*”. **(H)** Schematic drawings of representative patterns of guard cells and subsidiary cell clusters in *bdyda1-1*. **(I)** Gene/Protein diagram of *BdYDA1*. The vertical magenta bar indicates the *bdyda1-1* EMS mutation and the green bar indicates the *bdyda1-2* CRISPR/Cas9 induced mutation. Model generated in Gene Structure Display Server (Hu et al., 2015).

To determine whether the identified mutation was indeed causal for the phenotype, we generated *BdYDA1pro:BdYDA1-YFP:Yt*, a complementation construct consisting of ~5.1 kb of upstream sequence, the *BdYDA1* genomic region (including introns), a YFP tag, and ~1.5 kb of downstream sequence. The lateral organ and internode elongation defects of *bdyda1-1* were rescued by this construct (Fig. 2E), as were the stomatal patterning defects (Fig. 2C, quantified in 2F-G). This was strong evidence that the mutation in *BdYDA1* was responsible for the phenotype. Based on work *AtYDA* and other with MAPKKKs, mutations in the kinase domain could lead to hypomorphic or to null alleles (Sopeña-Torres et al., 2018; Lukowitz et al., 2004). We therefore created an additional clustered regularly interspaced short palindromic repeat (CRISPR)/CRISPR-associated protein 9 (Cas9) mutation-based allele in the first exon of *BdYDA1* to attempt to eliminate the protein altogether (Fig. S2A). *bdyda1-2* is heteroallelic for frameshift mutations predicted to encode truncated proteins of 41 and 95 amino acids in contrast to the normal BdYDA1 protein of 896 amino acids (Fig. S2C) and resulted in the same suite of stomatal and morphological phenotypes as *bdyda1-1*, however, the magnitude of the clustering was increased (Fig. 2D, S2B). The difference between the phenotypes produced by the E460>K missense allele compared to the truncation allele suggests that *bdyda1-1* is a hypomorphic allele. CRISPR/Cas9-induced mutations targeting the second *YDA* homologue [*BdYDA2* (Bradi3g51380)] did not result in any obvious stomatal or overall growth phenotypes (Fig. S3).

The large clusters of stomata in *Arabidopsis yda* mutants arise from aberrant asymmetric divisions in the self-renewing precursor cells before they commit to becoming GMCs (Bergmann et al., 2004). Such self-renewing asymmetric divisions, however, are absent in grasses. An advantage of the more streamlined and rigid developmental trajectory in grasses is that we could generate a pseudo-timecourse by imaging a single *Brachypodium* leaf from base to tip and observing the cells at different ontological stages in *bdyda1-1* and WT (Fig. 3A-H). To our surprise, early epidermal asymmetric divisions appeared normal in the *bdyda1-1* mutant (Fig. 3A). By the subsidiary cell recruitment stage (Fig. 3B), however, it was evident that the smaller daughters of the previous asymmetric division were not the only cells that had acquired stomatal fate. The larger daughter cells appeared to undergo extra divisions (Fig. 3B), and unusual subsidiary cell morphologies, or the spanning of two GC complexes by a single subsidiary cell (Fig. 3B-C) were consistent with supernumerary stomatal-row cells taking on GC identity. The later stages (Fig. 3C-D) of stomatal differentiation, including the symmetric GMC division and subsequent stomatal pore formation, occurred fairly normally, as they do in *atyda* mutants, suggesting that *BdYDA1* acts primarily in the early fate decisions and not in stomatal guard cell differentiation.

**Figure 3.**
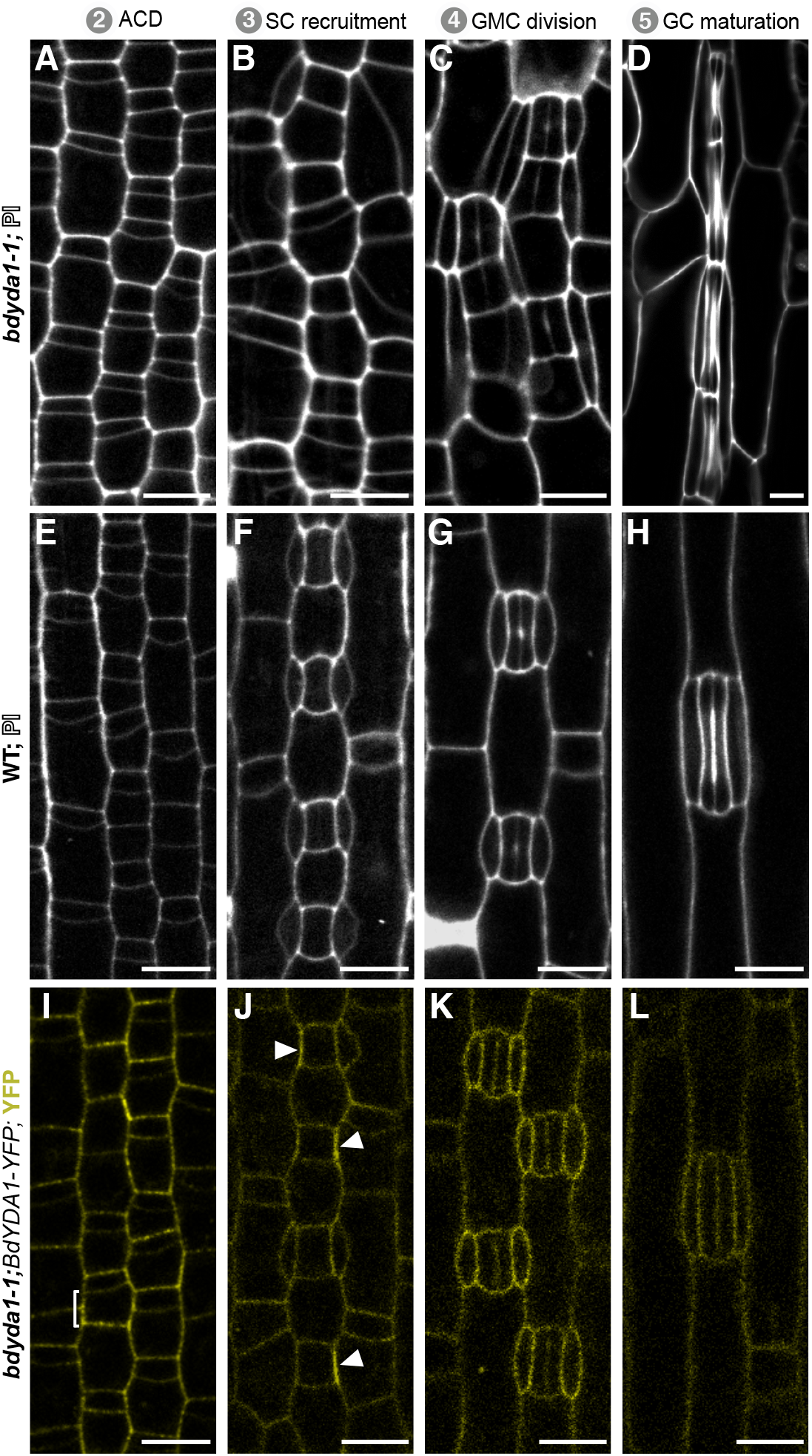
Imaging early development indicates that BdYDA1 is expressed throughout the stomatal lineage and that the initial defect in *bdyda1-1* appears to be improper enforcement of non-stomatal fates. Confocal images of progression of cells though four stages (as defined in Fig. 1A) of stomatal development in *bdyda1-1* mutants **(A-D)** and WT (Bd21-3) **(E-H)** [emerging 2^nd^ leaf at 6 dpg, stained with propidium iodide (PI)]. **(I-L)** Expression of rescuing *BdYDA1pro:BdYDA1-YFP:Yt* in *bdyda1-1* (T1 plant; emerging 2^nd^ leaf at 6 dpg; YFP channel only). Bracket in (J) and arrowheads in (K) indicate accumulations of transgene signal. Scale bar = 10 µm. All images are oriented with the base of the leaf blade (younger cells) towards the bottom and the tip of the leaf (older cells) towards the top.

To address how activity of BdYDA1 might enforce stomatal patterning in *Brachypodium*, we visualized the expression pattern of the complementing *BdYDA1pro:BdYDA1-YFP:Yt* reporter. BdYDA1-YFP signal was present in all stomatal lineage cells, from proliferative divisions until maturation (Fig. 3I-L); however, it was not specific to the stomatal lineage and appeared to be present at roughly comparable levels in all the different leaf epidermal cell types we monitored. Fluorescence was strongest in the cytoplasm and/or at the cell periphery and was not detectable in the nucleus. In some cells, BdYDA1-YFP fluorescence appeared to be concentrated at a single face of a cell, most frequently at the interface between a GMC and its neighbor cell that will give rise to a subsidiary cell (brackets in Fig. 3I and arrowheads in Fig. 3J; Fig. S4). This broad expression pattern is similar to that of AtYDA, suggesting that, like in *Arabidopsis*, it is the presence of appropriate upstream activating signals and downstream targets that provides specificity to YDA-mediated signaling activity.

### Cell fate marker expression suggests defects in fate reinforcement in *bdyda1-1*

To further dissect the defects during GMC specification and subsidiary cell recruitment observed in *bdyda1-1*, we examined the expression and behavior of stomatal lineage fate markers. *BdSCRM2pro:YFP-BdSCRM2* is a pan-stomatal lineage marker (Raissig et al., 2016). It can be visualized in nuclei from the time stomatal rows are specified; and, when these cells start dividing asymmetrically, BdSCRM2 is confined to the smaller daughter of these divisions (Fig. 4A) and remains restricted to stomatal complexes as they mature (Fig. 4B-D). In *bdyda1-1*, however, restriction of signal to the smaller daughter of an asymmetric division is lost (Fig. 4E-F). Specifically, we observed signal in larger cells between stomatal precursors (arrowheads in Fig. 4E). In some cases, these presumed pavement cell precursors underwent an ectopic asymmetric division generating a stomatal precursor positive for YFP-BdSCRM2 expression adjacent to the earlier-specified stomatal precursor (arrow in Fig. 4F).

**Figure 4.**
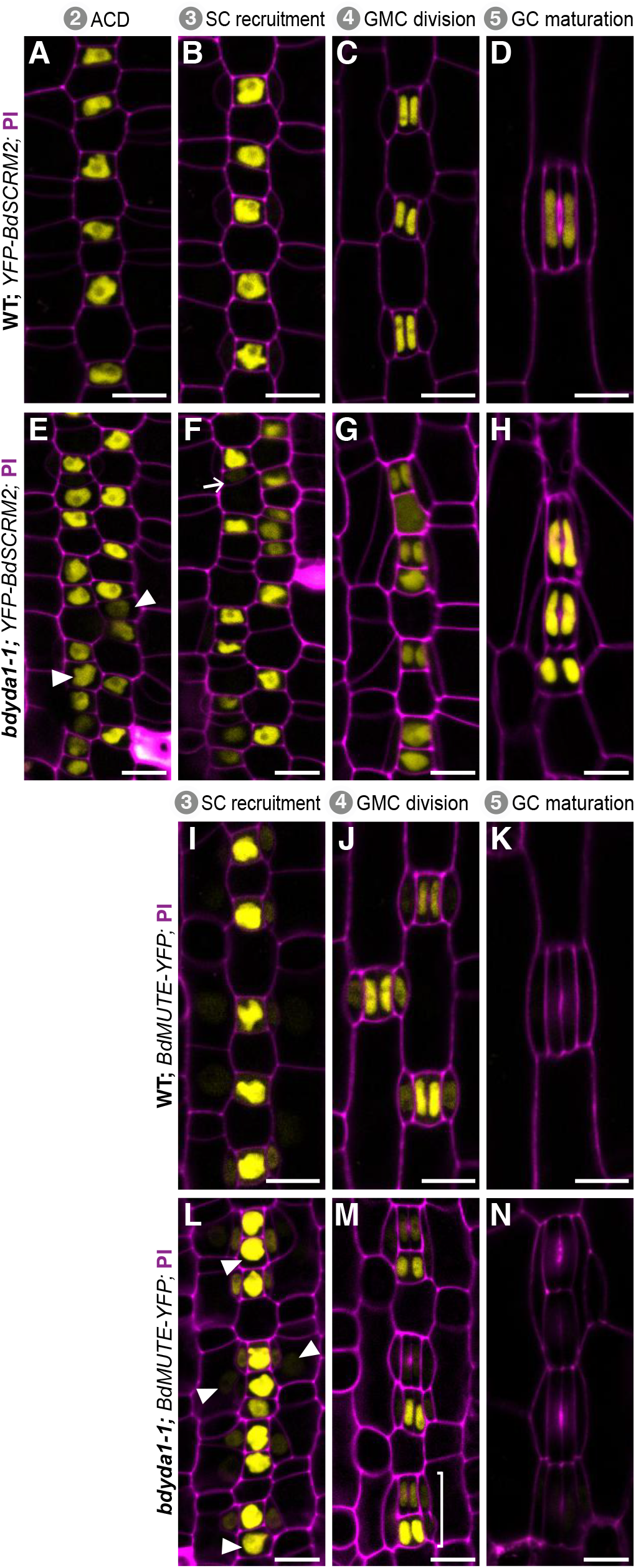
Misexpression of stomatal fate reporters in *bdyda1-1* mutants is consistent with the terminal fate specification defects. Confocal images of emerging 2^nd^ leaf of 6 dpg T1 plants. *BdSCRM2pro:YFP-BdSCRM2* reporter in WT (Bd21-3) **(A-D)** and *bdyda1-1* mutant **(E-H)** during stomatal development. Early in WT development, *BdSCRM2pro:YFP-BdSCRM2* appears only in the smaller daughter of an asymmetric division (A-B). However, at the same stage in the *bdyda1-1* mutant, signal is also present in mis-specified larger daughter cells (E-F). Arrowheads in (E) indicate examples of improper re-enforcement of non-stomatal fate in larger daughter of asymmetric division. Arrow in (F) points to an example of improper inhibition of division potential. *BdMUTEpro:BdMUTE-YFP* reporter in WT (**I-K)** and *bdyda1-1* mutant **(L-N)** during SC recruitment and GMC and SC specification. Reporter expression in WT is present only in GMCs, Subsidiary Mother Cells (SMCs), and SCs as stomata mature. In *bdyda1-1*, the same reporter also marks mis-specified and clustered GMCs, SMCs, and SCs. Arrowheads in (L) and bracket in (M) indicate ectopic marker expression during the SC recruitment and GMC division, respectively. Scale bar = 10 µm. Cell outlines are visualized with PI. All images are oriented with the base of the leaf (younger cells) towards the bottom and the tip of the leaf (older cells) towards the top.

To address later specification events and subsidiary cell recruitment, we analyzed the expression pattern of *BdMUTEpro:BdMUTE-YFP*, which, in WT, starts in young GMCs and shows strong signal in mature GMCs and weak signal in adjacent subsidiary mother cell files (Fig. 4I). BdMUTE expression is maintained until after GMC division in both GCs and subsidiary cells and disappears during complex maturation (Fig. 4J-K). In *bdyda1-1*, young and mature GMCs showed marked patterning defects, with numerous BdMUTE-positive cells positioned adjacent to one another (Fig. 4L). Marker expression was also present in subsidiary cells even if they were recruited and formed abnormally. BdMUTE persisted in stomatal clusters through GMC division, but was extinguished rapidly as the guard cells matured, as it was in WT (Fig. 4M-N).

Taken together, the reporter results agree with the morphological assessments and suggest that in *bdyda1-1*, mechanisms in place to control fate establishment and/or division potential early in the lineage are disrupted, but that stomatal differentiation and morphogenesis are largely unaffected.

### *BdYDA1* regulates cell patterning in other epidermal cell lineages

In studying the effects of *bdyda1-1* on the stomatal lineage, it became apparent that this was not the only epidermal cell lineage disrupted by the mutation. *Brachypodium* also produces regularly spaced hair cells in the leaf epidermis. These hairs arise via asymmetric divisions in a manner similar to stomata, and the two fates appear to be closely related and somewhat interchangeable [e.g., a cell file may contain a number of stomata, then a hair, then more stomata (Raissig et al., 2016); for an example, see Fig. 1B]. In *bdyda1-1* plants, the hair cell lineage displays defects very similar to the stomatal lineage, producing hair cells at higher density than in WT and often in longitudinal clusters (Fig. 5A-B, rescued in 5C, and quantified in 5D-E). In addition, the epidermis of the leaf sheath contains crenellated pavement cells and round silica cells sometimes accompanied by a small lens-shaped cell and a small triangular cell (Fig. 5F). Patterning and proliferation defects were evident in all of these cell types in *bdyda1-1*, and were complemented by the BdYDA1 reporter (Fig. 5G-H). We quantified patterning and proliferation defects in the large, round silica cells because, among the sheath cell types, they were most unambiguously identified. Like stomata and hair cells, the silica cells were produced at a greater density than in WT and were found in numerous longitudinal clusters (Fig. 5I-J).

**Figure 5.**
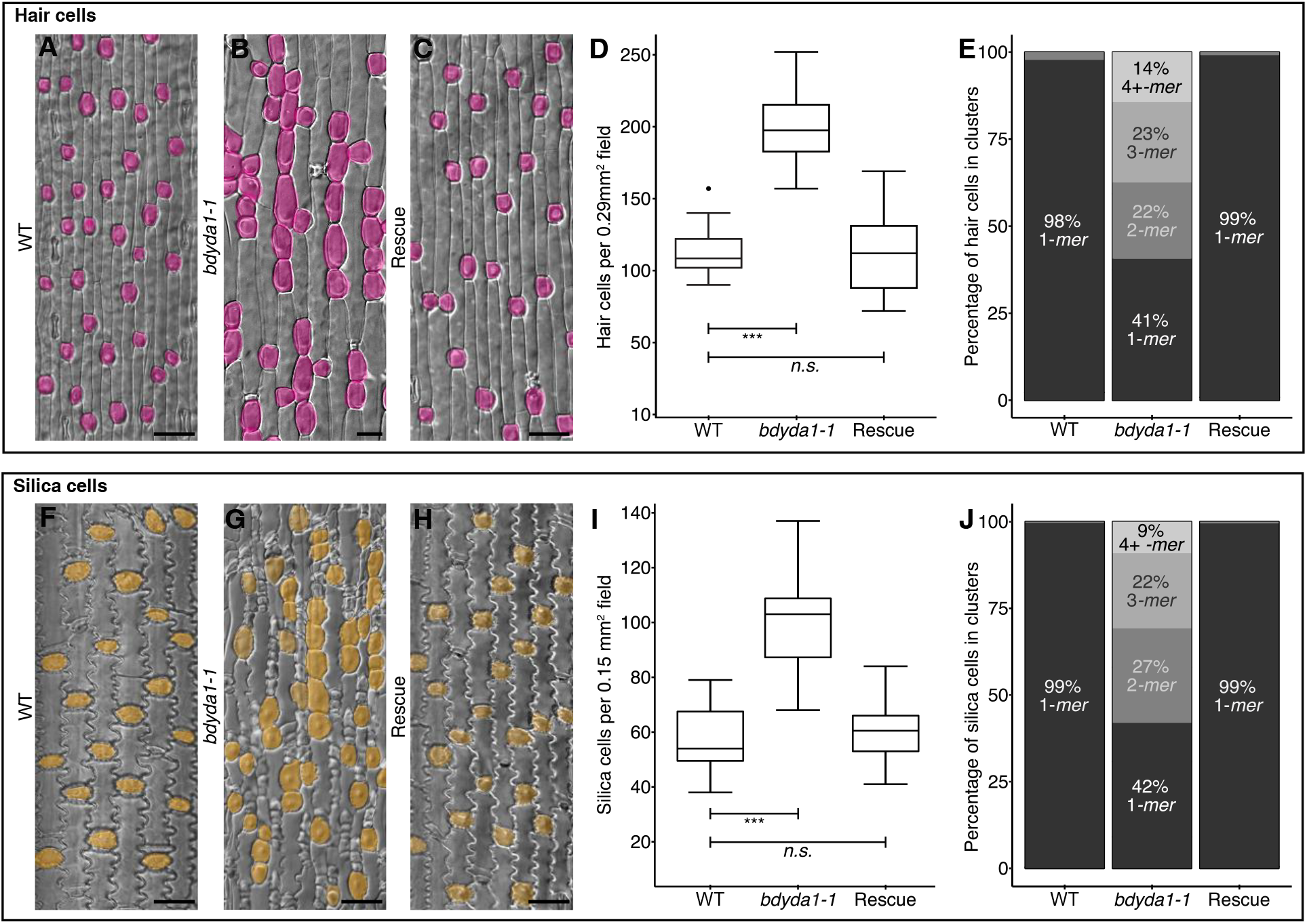
*bdyda1-1* mutants exhibit disruption of cell fates in other asymmetrically dividing epidermal lineages. **(A-C)** DIC images of cleared WT (Bd21-3) (A), *bdyda1-1* mutant (B), and *bdyda1-1* rescued with *BdYDA1pro:BdYDA1-YFP:Yt* (C) leaf epidermis. Hair cells are false-colored magenta. WT and complemented *bdyda1-1* images show 6^th^ leaf from base (3^rd^ from main tiller) 27 dpg. *bdyda1-1* images show epidermal peels of 6^th^ leaf from base (3^rd^ from main tiller) 27dpg. Scale bars = 40 µm. **(D)** Hair cell density of *bdyda1-1* mutants compared to that of WT and rescued *bdyda1-1* [6^th^ leaf from base (3^rd^ from main tiller) 27 dpg]. n=4 individuals for WT control and n=5 for rescued plants. For each sample, 5 different regions of the leaf were imaged and quantified. n=5 for *bdyda1-1* mutants for which 4 different regions of the leaf were peeled, imaged, and quantified. ***, significant at p<0.001; *n.s*., not significant (based on Kruskal-Wallis test followed by Dunn’s multiple comparison’s test). **(E)** Hair cell cluster profile as percentage of clustering of quantified hair cells in *bdyda1-1* mutants, WT, and rescued *bdyda1-1* (n=2286 hair cells for WT controls, n=3988 hair cells for rescue, and n=2789 hair cells for *bdyda1-1*). Clusters of 4 or more hair cells were grouped in last category “4+ *-mer*”. **(F-H)** DIC images of cleared WT (F), *bdyda1-1* mutant (G), and rescued *bdyda1-1* (H) sheath epidermis. Silica cells are false-colored orange. For all genotypes, images show the sheath of the 6^th^ leaf from base (3^rd^ from main tiller) 27dpg. Scale bars = 40 µm. **(I)** Silica cell density of *bdyda1-1* compared to that of WT and rescued *bdyda1-1* [sheath of the 6^th^ leaf from base (3^rd^ from main tiller) 27 dpg]. n=4 individuals for WT control, n=5 for rescued plants, and n=5 for *bdyda1-1* mutants. For all, 4-6 different regions of the sheath were imaged and quantified. ***, significant at p<0.001; *n.s.*, not significant (based on Kruskal-Wallis test followed by Dunn’s multiple comparison’s test). **(J)** Silica cell cluster profile as percentages of clustering of quantified silica cells in *bdyda1-1* mutants, WT, and rescued *bdyda1-1* (n=1328 silica cells for WT controls, n=2995 silica cells for rescue, and n=1691 silica cells for *bdyda1-1*). Clusters of 4 or more silica cells were grouped in last category “4+ *-mer*”.

## Discussion

By characterizing a mutation that disrupted stomatal patterning in *Brachypodium distachyon*, we identified the *BdYDA1* gene, a *Brachypodium* orthologue of the *Arabidopsis* MAPKKK gene *AtYDA*. Like *atyda* (Bergmann et al., 2004; Lukowitz et al., 2004), the *bdyda1* mutant produces excess stomata arranged in clusters and displays a stunted growth phenotype characterized by compressed internodes and compact lateral organs. Strikingly, we could show that clusters in *bdyda1* arise from misspecification of alternative fates in the epidermis, rather than due to alterations in the physical asymmetry of the divisions themselves. In addition, other non-stomatal epidermal cell types are also affected. Taken together, our results demonstrate that *BdYDA1* is a general regulator of cell fate establishment and enforcement, two processes that are crucial for the correct execution of asymmetric divisions during epidermal patterning of the grass leaf (Fig. 6A).

**Figure 6.**
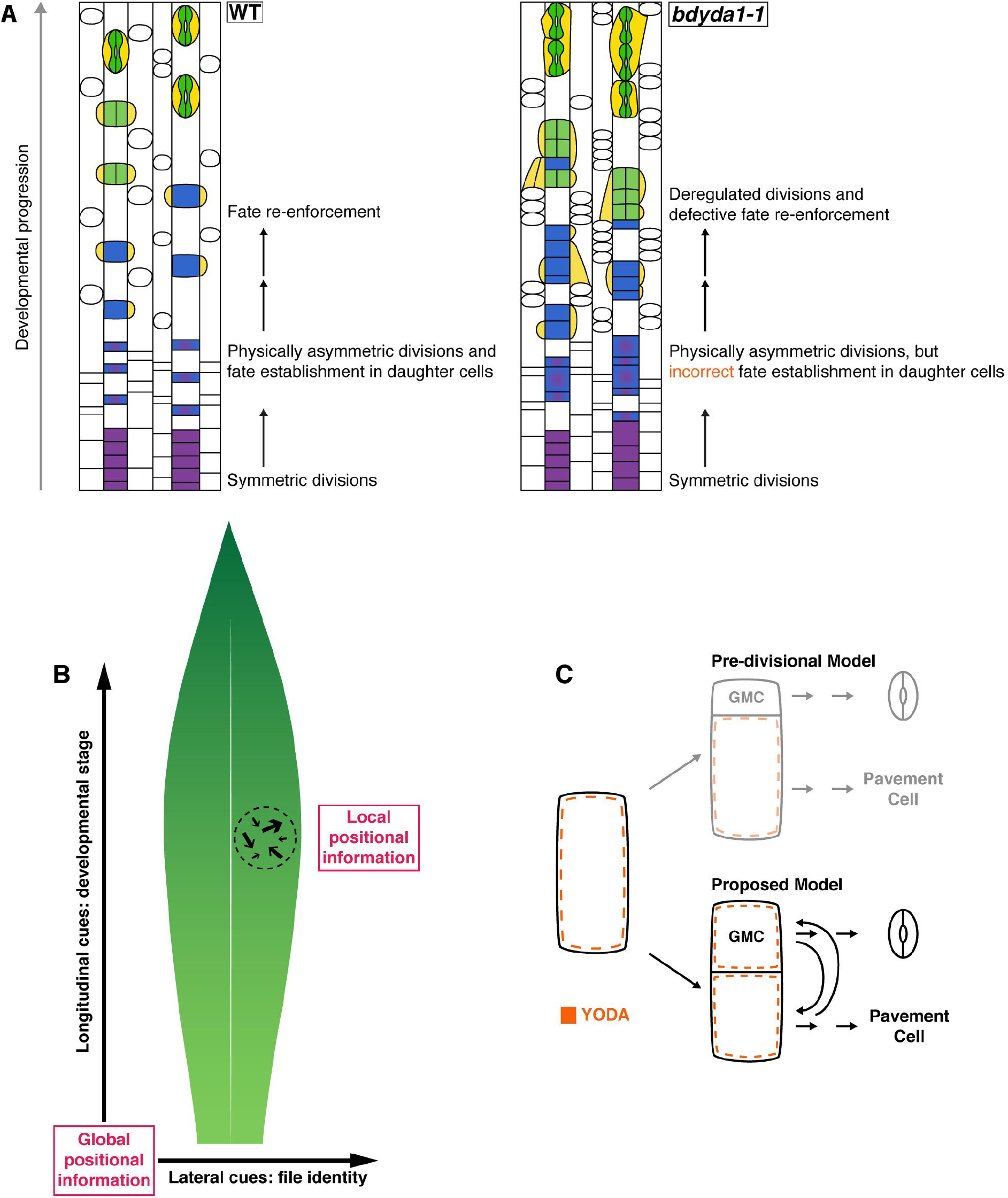
Summary of YDA’s proposed role in asymmetric divisions. **(A)** Schematic representation of the *bdyda1-1* phenotype and interpretations of the role of BdYDA1 in epidermal patterning of *Brachypodium* leaves. **(B)** Global and local positional information feed into developmental decisions that orient and position stomatal precursors. Global positional information in the form of lateral and longitudinal cues direct cell file identities and developmental progression of lineages, respectively. Local positional information controls fate re-enforcement to establish the correct pattern and distribution of stomata and their precursors. The relative influence of global vs. local sources of positional information is likely to be species-specific, i.e. longitudinally growing grass leaves are more heavily influenced by global cues and radially growing leaves with self-renewing stem-like divisions, like *Arabidopsis*, by local cues. **(C)** In contrast to *Arabidopsis*-derived pre-divisional models which suggested YDA mainly acts to establish physical asymmetry prior to fate establishment, we propose that YDA is primarily a post-divisional fate re-enforcer. This requires that YDA be present in both daughters of an asymmetric cell division and be available for reciprocal (and continuous) signal transduction downstream of cell-cell communication systems.

Asymmetric divisions are produced and oriented by a combination of intrinsic and extrinsic cues. AtYDA plays roles in the transduction of both sources of developmental information in *Arabidopsis* stomatal development. Positional (extrinsic) information conveyed by EPFL family ligands interacting with ERf/TMM receptors can activate the AtYDA MAPK cascade, leading to AtSPCH inhibition and repression of stomatal identity (reviewed in Pillitteri and Torii, 2012). More recently, a role in intrinsic polarity via a physical association between AtYDA and the polarity factor BASL emerged (Zhang et al., 2015). The cortical AtYDA/BASL complex is preferentially inherited by the larger daughter cell of an asymmetric division, leading to differential signaling capacity; the cell with higher AtYDA has lower AtSPCH levels and consequently loses stomatal identity (Zhang et al., 2016).

Could BdYDA1 participate in similar signaling or intrinsic polarity modules? The observation of non-uniform BdYDA1-YFP distribution around cells could be a hint to its participation in a differentially inherited polarity complex, but if this were the case, it would require an as-yet unknown polarity partner, since *BASL* homologues have not been detected in the grasses. Moreover, the specific location of the BdYDA1-YFP enrichment, at the boundary between GC and prospective SC, is not consistent with it being preferentially segregated to the larger (non-stomatal) cell during normal development. In terms of the cell-cell signaling response, *Arabidopsis* YDA and *Brachypodium* YDA1 proteins show appreciable sequence similarity (65% overall and 90% in kinase domain; Fig. S5) and components of the YDA MAPK pathway have clear orthologues in *Brachypodium*, including downstream kinases *MKK4* (Bradi1g46880), *MKK5* (Bradi3g53650), *MPK3* (Bradi3g53650), and *MPK6* (Bradi1g49100). Upstream signaling components include multiple EPFL family members, *TMM* (Bradi2g43940), and *ERECTA*, although the *ERECTA* family consist of only two members (Bradi1g46450 and Bradi1g49950).

While the presence of homologous signaling pathway genes makes participation in parallel signaling cascades possible for AtYDA and BdYDA1, there remains the issue of the distinct ontogeny of dicot and grass stomata. With no self-renewing divisions, mature complexes restricted to specific cell files, and consistent orientation of each complex along the leaf’s proximal-distal axis, the positional information required in grasses is very different from that in *Arabidopsis* where distributed “point sources” of signals and extensive neighbor to neighbor signaling are dominant patterning mechanisms (Torii, 2015) (Fig. 6B). As disorderly as epidermal identities become within *bdyda1* cell files, they still obey tissue-wide fate arrangements, indicating that different factors control lateral (and proximal-distal) positional information in the leaf. While none of the work with grass EPF homologues has yet demonstrated an effect on lateral patterning (Hughes et al., 2017; Yin et al., 2017), expanded expression of SHORTROOT, a factor normally involved in internal cell fates, leads to production of supernumerary stomatal rows in rice (Schuler et al., 2018) suggesting that lateral information may be provided through different pathways.

When considering downstream targets of a BdYDA1-mediated MAPK cascade in cell fate reinforcement roles, the stomatal initiation module could be targeted to inhibit stomatal fate establishment in larger daughter cells of asymmetric divisions, much like it is in *Arabidopsis*, but the precise protein target in this complex may be different. In *Arabidopsis*, AtSPCH is the demonstrated target of MAPK regulation (Lampard et al., 2009), but in *Brachypodium*, BdICE1 may play a more dominant role since it possesses predicted high-fidelity MAPK target sites within the protein degradation-associated PEST domain, while BdSPCH1/2 do not (Raissig et al., 2016). Furthermore, expression of ubiquitin-promoter driven YFP-BdICE1 accumulates only in the stomatal lineage cells of the leaf, indicating that this protein is subject to posttranslational regulation, as one would expect from a target of a MAPK cascade (Raissig et al., 2016).

BdYDA1 also regulates fate re-enforcement in other non-stomatal epidermal cell linages; however, by what means it does so remains to be explored. It is likely that BdYDA1 has targets that could enforce fate asymmetry in all of these decisions, while the specific fate of the cells (stomata, hair, or silica) would be determined by other information. Such is the case with *Arabidopsis* embryos and stomata where signaling can work when swapped between these developmental contexts (Bayer et al., 2009), but downstream transcription factors provide unique cell identities (Jeong et al., 2011; Ueda et al., 2017). We noticed that in the early truncation allele *bdyda1-2* the degree of clustering is less in non-stomatal epidermal cells than in stomatal files (Fig. 2D and S2B) suggesting that BdYDA1- independent fate determining mechanisms also exist in these lineages.

In *Arabidopsis* embryos, roots, and stomatal lineage, loss or overactivity of *AtYDA* changes an asymmetric division into one whose daughters exhibit equalized cell fates and cell sizes. From these phenotypes, it is intuitive to imagine AtYDA’s role as one initiating asymmetry in the mother cell of these formative divisions (Fig. 6C). Furthermore, considering that YDA’s downstream effectors are the microtubule regulating kinases AtMPK3/6, AtYDA was suggested to mediate cytoskeletal behaviors leading to the placement of division planes and creation of asymmetric divisions (Smékalová et al., 2014). In our present study, however, we showed that the physical asymmetry of divisions is not affected in the absence of *BdYDA1*, prompting us to re-evaluate whether AtYDA actually generates pre-divisional, physical asymmetry, or whether the failure to create different-sized cells in *atyda* mutants stems from a post-divisional failure in cell identity. High-resolution time-lapse imaging of developmental decisions in *Arabidopsis* would be needed to distinguish between *YDA* playing primarily a pre- or post-divisional role.

The work presented here emphasizes the value of comparative developmental studies, as the *bdyda1* mutations reveal roles for the YDA pathway in numerous cell-fate decisions, some of which represent cell types not present in *Arabidopsis*. This work also provokes a conceptual shift in our emphasis on MAPK signaling as required for the creation of asymmetry to MAPK signaling required for the post-divisional enforcement of asymmetric fates. In a kingdom devoid of Notch-Delta lateral inhibition systems with scant evidence for any segregated fate determinants, and one characterized by exceedingly flexible development, it may be that plant cell fate decisions are rarely made in advance, but are subject to multiple rounds of confirmation through post-divisional communication and re-enforcement.

## Materials and Methods

### Plant Material

The *bdyda1-1* mutant was recovered from the M3 generation of an ethyl methanesulphonate (EMS) mutagenesis of Bd21-3 ecotype (seeds provided by Dr. John Vogel, JGI; Raissig et al., 2016), and maintained through selection of heterozygous individuals. Bd21-3 was used as WT for all experiments described in this paper (Vogel and Hill, 2007).

Plants were initially grown on ½ strength MS agar plates in a 22ºC chamber with a 16-hour light/8-hour dark cycle (110 µmol m^−2^ s^−1^), then subsequently transferred to soil and placed in a greenhouse with a 20-hour light/4-hour dark cycle (250-300 µmol m^−2^ s^−1^; day temperature = 28ºC, night temperature = 18ºC). Seeds were stratified for at least two days at 4ºC before transfer to light.

### Generation of Constructs

*BdSCRM2:YFP-BdSCRM2* was described in Raissig et al., 2016. To create *BdYDA1pro:BdYDA1-YFP:Yt* and *BdMUTEpro:BdMUTE-YFP*, sequences were amplified from BACs BD_ABa0027F22 and BD_ABa0042O15/BD_AB0008G12 (Arizona Genomics Institute; http://www.genome.arizona.edu/), respectively. For *BdYDA1pro:BdYDA1-YFP-Yt*, 5.1 kb upstream sequence (primers *BdYDA1*pro5.1kb-1F and *BdYDA1*pro_AscI-1R) and 3’ sequence (*BdYDA1*term-1F and *BdYDA1*term-1R) of the *BdYDA1* gene were cloned into pIPKb001 (Himmelbach et al., 2007). This then was recombined with a *BdYDA1*-*YFP* fusion in pENTR-D, composed of *BdYDA1* genomic region (primers *BdYDA*1proPacI-1F and *BdYDA1*cDNAnscAscI-1R) followed by AscI-flanked *Citrine* YFP.

For *BdMUTEpro:BdMUTE-YFP*, 1.1 kb upstream sequence of the *BdMUTE* gene (Bradi1g18400) (primers *BdMUTE*pro-FWD and *BdMUTE*pro-REV) was cloned into pIPKb001t (Raissig et al., 2016). Separately, the *BdMUTE* genomic sequence (primers *BdMUTE*-CDS-FWD and *BdMUTE*-CDS-REV) was cloned into pENTR-D with a poly- alanine linker (annealed primers Ala_linker-F and Ala_linker-R) and an AscI-flanked *Citrine* YFP inserted 3’ of the gene by AscI digest. Finally, the entry clone was recombined into the pIPKb001t vector.

CRISPR constructs were designed using the vectors pH-Ubi-cas9-7 and pOs-sgRNA (vectors and protocol described in Miao et al., 2013). The online server, CRISPR-P, was used to identify candidate spacer sequences (Lei et al., 2014). Spacers were generated by annealing oligo duplexes priMXA38F+39R for *BdYDA1* CRISPR_sgRNA_6 (which generated *bdyda1-2)* and priMXA30F+31R for *BdYDA2* CRISPR_sgRNA_11. Primers priMXA50 and 52 were used to genotype *bdyda1-2* and primers priMXA48 and 49 were used to genotype *bdyda2-1* and *bdyda2-2*.

### Generation of transgenic lines

*Brachypodium* calli derived from Bd21-3 and *bdyda1-1*/+ parental plants were transformed with AGL1 *Agrobacterium tumefaciens*, selected and regenerated according to standard protocols (http://jgi.doe.gov/our-science/scienceprograms/plant-genomics/brachypodium/). Plants and calli from *bdyda1-1*/+ parents were genotyped using primers priMXA25, 26, and 27 in a single PCR reaction as described in (Gaudet et al., 2009).

### Microscopy and Phenotypic Analysis

For imaging on a Leica SP5 confocal microscope, leaves were carefully taken out from the surrounding sheath and stained in propidium iodide (1:100 dilution of 1 mg/ml stock) to visualize cell walls, then mounted in water. For DIC imaging on a Leica DM6 B microscope of WT and rescue plant leaf (distal 1.5 to 2 cm of the 6^th^ leaf blade of 22dpg leaves) and sheath (top 1 cm of the 6^th^ leaf sheath of 22dpg plants) tissue were collected into 7:1 ethanol:acetic acid and incubated overnight to remove chlorophyll, then rinsed with water, and mounted in Hoyer’s medium to clear. The same was done for DIC imaging of *bdyda1-2*, except no sheath tissue was collected for it. For DIC imaging of *bdyda1-1*, epidermal peels were collected from the tissue of interest into Hoyer’s medium, mounted on slides, and examined. Cell numbers and extent of stomatal clusters were counted directly on a computer display attached to the microscope (0.29mm^2^ field of view for stomata and hair cell counts; 0.15mm^2^ field of view for silica cell counts). Only cells fully contained within the image frame were included in the respective analysis. In cases when a cluster expanded beyond the image frame, cells outside the image frame were included to correctly represent the number of cells part of the cluster. For SEM images, WT leaves were taken directly from a growing plant, then introduced into an Environmental SEM (FEI Quanta 200) without any further treatments.

### Statistical Analysis and Plotting

Statistical analysis was performed in R. The Shapiro-Wilk test (shapiro.test function) was used to check samples for normality; as many samples displayed non-normal distributions, the Kruskal-Wallis test (a nonparametric analogue of ANOVA; kruskal.test function) was used, followed by Dunn’s Multiple Comparisons tests (dunn.test, dunnTest) to assess significance of pairwise comparisons.

## Acknowledgments

We thank Akhila Bettadapur for technical assistance, Dr. John Vogel (JGI) for creating *Brachypodium* resources, Dr. Kathryn Barton (Carnegie, DPB) for use of her SEM, and Dr. Michael Raissig and other members of the Bergmann lab for discussions and comments on the manuscript.

## Contributions

EA and MXAG performed all experiments except creation of BdMUTE reporter, done by JM. MXAG created figures.

EA and MXAG did quantifications and statistical analysis.

EA, MXAG, and DCB designed experiments and wrote the manuscript.

